# Assessing the longevity of the Theta-induced Memory Effect (TIME)

**DOI:** 10.64898/2026.07.20.739535

**Authors:** Eleonora Marcantoni, Danying Wang, Kimron L. Shapiro, Simon Hanslmayr

**Author notes:** Corresponding authors: Eleonora Marcantoni, Simon Hanslmayr.

## Abstract

Theta-phase synchronisation between visual and auditory rhythmically modulated sensory stimuli during encoding has been shown to enhance associative memory, an effect known as Theta-Induced Memory Effect (TIME). However, whether this advantage is short-lived or persists over time remains unknown.

To address this, we examined the TIME effect in 52 participants randomly assigned to a short-delay group (immediate retrieval) or a long-delay group (retrieval after 24 hours). Electroencephalography (EEG) was recorded during incidental encoding of audio-video pairs. The luminance of the video clips and the amplitude of the sounds were modulated at theta frequency (4 Hz) and presented either in-synchrony (0° phase offset between audio and video) or out-of-synchrony (the sound 180° phase-shifted relative to the video). Source reconstruction was used to estimate the actual phase difference between visual and auditory cortices for every trial.

At the behavioural level, no significant difference in memory accuracy was found between the synchronous and asynchronous conditions in either group. However, after reconstructing the actual phase difference at the source level, the TIME effect was replicated in the short-delay group, with significantly higher memory accuracy at the optimal reconstructed theta phase compared to out-of-phase. In contrast, the long-delay group showed a reversed pattern, with higher accuracy in the asynchronous condition, though this difference did not reach significance. These findings suggest that the TIME effect is time-limited and may not survive or even reverse after overnight consolidation.

## 1 Introduction

Episodic memory allows us to remember rich multisensory experiences from our lives. The hippocampus is thought to play a central role in forming these memories by binding information from different sensory modalities into a unified representation (Moscovitch, 2008; Scoville & Milner, 2000). At the cellular level, associative memory formation is thought to depend on Hebbian synaptic plasticity. According to this framework, connections between co-active neurons strengthen through long-term potentiation (LTP; Bliss & Lømo, 1973). Evidence from rodent literature suggests that whether LTP or long-term depression (LTD) occurs at hippocampal synapses depends critically on the phase of the ongoing theta rhythm at which stimulation is delivered, with stimulation at the peak of theta inducing LTP and stimulation at the trough inducing LTD (Hölscher et al., 1997; Huerta & Lisman, 1995; Hyman et al., 2003). These findings suggest that the relative timing of neural activity may influence whether information becomes successfully bound into memory.

To test this idea in humans, our group developed a rhythmic sensory stimulation (RSS) paradigm to directly manipulate the degree of synchrony between visual and auditory stimuli to be encoded during an associative memory task (Clouter et al., 2017). In this paradigm, the luminance of video clips and the amplitude of sound clips are rhythmically modulated at theta frequency (4 Hz). Critically, the two streams are presented either in-synchrony or out-of-synchrony. The rationale is that synchronous streams are more likely to arrive at the hippocampus simultaneously, increasing the probability of LTP-dependent binding compared to asynchronous inputs. Consistent with this prediction, we found better associative memory for in-phase than out-of-phase audiovisual pairs. This effect was specific to theta frequency and was not observed at other stimulation frequencies, hereafter referred to as the Theta-Induced Memory Effect (TIME).

Subsequent work has provided further support for the TIME effect. In a following study, we replicated the behavioural finding and used EEG to show that memory performance was related to the trial-by-trial phase difference between entrained activity in visual and auditory cortices (Wang et al., 2018). Extending the TIME effect beyond declarative memory, Plog et al. (2022) demonstrated that theta-phase synchronisation between a conditioned stimulus and an unconditioned stimulus facilitated fear memory formation, then replicated online in a larger sample (Plog et al., 2023). More recently, Biau et al. (2025) showed that in an audiovisual speech memory task, theta oscillations in the neocortex and hippocampus were stronger for synchronous than asynchronous audiovisual speech clips during encoding, and that asynchrony between lip movements and auditory speech reduced subsequent memory reinstatement.

However, not all replication attempts have been successful. Van der Plas et al. (2020) found no evidence for enhanced memory when theta synchronisation between visual and auditory cortex was induced via transcranial alternating current stimulation (tACS) of the visual cortex. Simeonov and Das (2025) similarly failed to replicate the effect in an online study involving emotive stimuli. In addition, both Serin et al. (2024) and our group (Wang et al., 2026) attempted to replicate the TIME effect using MEG, and neither replicated the significant effect when memory performance was averaged across trials. Notably, in Wang et al. (2026), we did observe the effect at the single-trial level when trials were re-labelled according to the actual phase difference reconstructed from the sensory cortices. Furthermore, pre-stimulus alpha power was found to affect the strength of entrainment of these cortices, suggesting that the TIME effect is brain-state dependent.

These mixed findings point to the importance of better understanding the boundary conditions of the TIME effect. Beyond theoretical interest, RSS is a non-invasive, easy to implement, and low-cost technique with potential therapeutic applications for populations with memory impairments. Realising this potential requires a deeper understanding of when and why the TIME effect occurs, and how long the effect persists.

One important yet unexplored aspect of the TIME effect is its temporal longevity. Whereas LTP- like mechanisms have been proposed to account for the TIME initial memory advantage, it remains unclear how this encoding advantage is affected by subsequent offline consolidation processes. Recent findings support the view that this synchrony-related memory advantage induced by theta RSS is mediated by hippocampal associative binding (Biau et al., 2025; Marcantoni et al., 2026, preprint). If so, understanding the long-term consequences of the TIME effect requires considering how hippocampus-dependent memories evolve after encoding. If theta RSS facilitates hippocampal associative binding through LTP-like mechanisms, then in-phase audiovisual pairs may form stronger hippocampus-dependent representations. According to Active Systems Consolidation theory (Diekelmann & Born, 2010; Klinzing et al., 2019; Rasch & Born, 2013) newly acquired hippocampus-dependent memories are actively reorganised during offline consolidation through repeated hippocampal replay and their gradual integration into distributed neocortical networks. If in-phase stimulation produces a stronger initial hippocampal memory trace, the relative advantage over out-of-phase pairs may persist, or even become more robust, after an overnight delay. At the same time, sleep-dependent consolidation is thought to involve both selective replay and global synaptic renormalisation. The Synaptic Homeostasis Hypothesis (SHY; Tononi & Cirelli, 2006, 2014) proposes that synaptic potentiation accumulated during wakefulness is downscaled during sleep, restoring synaptic efficiency and preventing saturation. From this perspective, an encoding-related advantage produced by synchronous stimulation may be reduced after overnight sleep because global renormalisation compresses the overall distribution of synaptic weights, reducing the relative difference between initially stronger and weaker memory traces.

The aim of the present study was to address this question by examining the TIME effect at two retrieval delays: immediately after encoding (short-delay group) and approximately 24 hours later following overnight consolidation (long-delay group). We predicted that the TIME effect would be present in the short-delay group, replicating previous findings. For the long-delay group, we considered two competing possibilities: the synchrony-related memory advantage could persist, or it could be reduced or absent after overnight consolidation.

## 2 Methods

### 2.1 Sample

Participants were recruited from the University of Glasgow School of Psychology and Neuroscience’s online subject pool and screened prior to testing. Inclusion criteria required normal or corrected-to-normal vision, normal hearing, and no contraindications to RSS, including no history of photosensitive epilepsy or migraine. The experiment was performed in accordance with the ethical standards of the Declaration of Helsinki and received approval from the local ethics committee. All participants gave written informed consent and were compensated for their participation. A total of 63 healthy participants aged 18–36 years took part in the study. Data from 11 participants were excluded from the analyses due to performance at or below chance level. To determine the chance threshold, we used an exact binomial test (chance = 0.25, given four response alternatives; one-tailed α = .05), which resulted in a minimum of 13 correct responses out of 32 trials to be considered above chance (accuracy ≥ .41). The remaining 52 participants (short-delay group: n = 24, n females = 12, mean age ± SD = 23.2 ± 3.78 years; long-delay group: n = 28, n females = 14, mean age ± SD = 24.4 ± 4.32 years) were included in all subsequent analyses. All participants were right-handed except for 4 left-handed participants (short group = 3; long group = 1).

### 2.2 Stimuli

Visual and auditory stimuli were taken from the same stimulus pool as those used in Clouter et al. (2017) and Wang et al. (2018). Movie clips of 3 s duration were presented at 30 frames/s, resulting in 91 frames per clip. The movie clips depicted natural scenes, animals, architecture, or human activities, and were selected such that the visual and auditory content was not semantically related. Luminance was modulated sinusoidally at 4 Hz between 0% and 100%, with all clips initially starting at 50% luminance. Sound amplitude was modulated sinusoidally at 4 Hz between 0% and 100%, with a sine wave at either 0° or 180° phase offset relative to the luminance modulation of the movie clips. Each sound clip was drawn from one of four sound categories (acoustic choir, movie soundtracks, and orchestra), as described in Wang et al. (2018). Each sound clip was presented concurrently with a movie clip for 3 s, with an onset lag of 40 ms relative to the movie. This lag compensated for the faster processing of auditory relative to visual stimuli and ensured that the targeted phase difference of 0° or 180° was achieved between auditory and visual cortex. The assignment of phase offset conditions (0° or 180°) to audio-video pairs was counterbalanced across participants, with the constraint that each participant from the short-delay group was matched with a participant from the long-delay group who received the same stimulus set.

Stimuli presentation was controlled using the Psychophysics Toolbox extensions (Brainard, 1997; Kleiner et al., 2007; Pelli, 1997) in MATLAB (R2022a; MathWorks, Natick, MA, USA). Visual stimuli were displayed on a VIEWPixx /3D Lite LCD monitor (VPixx Technologies Inc., Saint-Bruno, QC, Canada) with a refresh rate of 120 Hz. Auditory stimuli were delivered binaurally via insert earphones (ER-3C, Etymotic Research Inc., IL, USA). The phase offsets between audio and video in the 0° and 180° conditions were checked before the study started using the Black Box ToolKit (BBTK v3; The Black Box ToolKit Ltd., England, UK). Stimuli onset triggers were delivered to the EEG recording system via the VIEWPixx pixel mode, in which a pixel in the corner of the display changes colour to generate a TTL pulse. The trigger was configured such that the pixel colour change coincided with the display of the first frame of each video clip.

### 2.3 Procedure

Participants were randomly assigned to one of two experimental groups differing in retrieval delay: a short-delay group, in which memory was tested immediately after encoding, and a long-delay group, in which memory was tested approximately 24 hours later. The experimental session consisted of three tasks in the following order: a unimodal localiser task, an associative memory task, and a synchronisation task. The associative memory task comprised three phases: audio-video pair ratings, a distractor, and a retrieval phase. For participants in the long-delay group, the retrieval phase was completed in a second session on the following day, during which EEG was not recorded. Throughout the session, participants were seated in a dimly lit sound-attenuated booth, with their head stabilised using a chin rest, approximately 60 cm from the screen.

Participants were not informed that their memory for the audio-video pairs would be tested to prevent the use of intentional encoding strategies. Instead, they were told that the study investigated how neural oscillations relate to performance across several cognitive tasks. Consequently, participants completed the initial audio-video ratings without any expectation that their memory for the associations would later be assessed. Participants assigned to the long-delay group were additionally informed that these tasks would be completed across two sessions to minimise fatigue. All participants assigned to the long-delay group returned for the second session.

#### 2.3.1 Unimodal localiser task

To localise auditory cortical sources in the EEG source reconstruction, participants completed a unimodal auditory task at the beginning of the experimental session. In the previous studies, the unimodal task was administered after the main experiment and used modulated sound clips similar to those used in the associative memory task. In the present study, the unimodal task was moved to the beginning of the session because the long-delay group did not wear the EEG cap during the second session, i.e., the unimodal task had to be completed during the first session. Therefore, to avoid interference with the experimental stimuli to be encoded in the subsequent associative memory task, pure tones amplitude-modulated at 4 Hz were used instead. The pure tones corresponded to musical-note frequencies spanning C3 to B5 (130.81-987.77 Hz) and were presented for 3 s with jittered inter-trial intervals (1-3 s). On each trial, participants rated how pleasant they found the sound on a scale from 1 (very unpleasant) to 5 (very pleasant). The task comprised 50 trials, consistent with previous experiments. Visual stimuli were not included, as visual source localisation could be reliably estimated from the audio-video pairs presented during the associative memory task (further explained in Section 2.5.2).

#### 2.3.2 Associative memory task

The associative memory task consisted of three phases: ratings, distractor, and retrieval. During the ratings phase, participants viewed 32 unique audio-video pairs and rated how well the audio suited the video on a scale from 1 (not at all) to 5 (very well). On each trial, a fixation cross was displayed at the centre of the screen for a jittered interval ranging from 1 to 3 s, followed by the 3-second pair presentation, after which participants provided their rating response. Each audio-video pair was repeated 4 times across blocks to help memorisation despite participants being unaware of the subsequent memory test. This resulted in a total of 128 trials, divided into 8 blocks of 16 trials each.

Following the ratings, participants completed a 30-second distractor task in which participants were asked to count backward in steps of 3 out loud, starting from a random number between 170 and 199 that was displayed on the screen. This was designed to minimise short-term memory contributions and ensure that subsequent performance reflected long-term associative memory.

Finally, in the retrieval phase, memory was tested using a four-alternative forced-choice recognition task (i.e., a chance level of 25%). Participants were presented with each of the 32 sound clips together with four video still frames displayed simultaneously on the screen. They were instructed to select the video that had been paired with the sound during the ratings phase. The 32 retrieval trials were presented in 2 blocks of 16 trials.

#### 2.3.3 Synchronization task

After the associative memory task, participants completed a synchronisation task in which they judged whether the audio and video of 16 pairs were presented in synchrony or out of synchrony. Trials consisted of a jittered fixation period (1-3 s) followed by a 3-s presentation of an audiovisual pair. This task was included to verify that participants could not perceptually discriminate between the 0° and 180° phase offset conditions.

### 2.4 Acquisition and preprocessing

EEG was recorded using a BioSemi ActiveTwo system (BioSemi B.V., Amsterdam, Netherlands) with 128 scalp electrodes at a sampling rate of 2048 Hz. Data were recorded using the ActiView software. Electrode offsets were monitored online via the ActiView interface and kept within acceptable ranges. Electrode positions were digitized using a Polhemus Space3 Fastrak device (Polhemus, Colchester, VT, USA) within Brainstorm (Tadel et al., 2011); version 3.230301) running in MATLAB (R2022b).

Prior to EEG preprocessing, digitized electrode positions were visually inspected to identify any inaccuracies. EEG preprocessing was performed using FieldTrip (Oostenveld et al., 2011); version 20231025) running in MATLAB (R2023b). Continuous data were bandpass filtered between 0.5 and 80 Hz (third-order Butterworth filter), and a bandstop filter was applied at 49–51 Hz to remove line noise. Data were then segmented into epochs of −2000 to 5000 ms relative to stimulus onset and resampled to 512 Hz. Bad channels and trials containing gross artefacts were identified and removed through visual inspection. Independent Component Analysis (ICA) was subsequently performed using the runica algorithm. Components were identified as artefactual based on their scalp topographies and time courses, targeting eye blinks, horizontal eye movements, and cardiac activity. After removal of these components, the remaining components were back-projected to reconstruct the EEG signals. The reconstructed data were visually inspected to confirm adequate removal of the targeted artefacts. Bad channels identified during the initial artifact rejection were interpolated using a weighted triangulation method based on neighbouring electrodes. Data were then re-referenced to the average of all channels. A final round of visual inspection was performed to remove any remaining artefacts.

### 2.5 EEG Source reconstruction

Although the phase offset between the visual and auditory stimuli was experimentally controlled, the phase difference between visual and auditory cortices does not necessarily equal the experimentally defined phase offset, due to variability in the transduction delays and in the brain state of the participant at the time of stimulation (Wang et al., 2026). To account for this trial-by-trial variability, source reconstruction was performed to estimate the actual phase difference between the visual and auditory cortices at the time of encoding for each trial. This analysis was performed using a pipeline developed by our group (Wang et al., 2018), recently refactored and made openly available (Hainke et al., 2026, preprint; see the Data and Code Availability section for access to the source code). The pipeline consists of four main stages: source space construction, unimodal source reconstruction, multimodal source reconstruction, and trial-level phase back-sorting.

#### 2.5.1 Source space construction

To define the source space for subsequent beamforming analyses, subject-specific headmodels and leadfields were constructed for each participant. In the absence of individual MRI scans, a template boundary element model (BEM) from FieldTrip was used for all participants. Each participant’s digitized electrode positions were aligned to the template headmodel through a three-step procedure: fiducial-based alignment using the nasion and left and right preauricular points, followed by interactive manual refinement, and finally projection of the electrodes onto the head surface. A subject-specific leadfield was then computed on a regular 10 mm grid spanning the brain volume.

#### 2.5.2 Unimodal source localization

The first step was to identify the location of peak neural activity in response to the 4-Hz modulated auditory and visual stimuli. This enables the identification of source-specific spatial filters that can subsequently be applied to the multimodal data to obtain clean, separated trial-level time series for each ROI. Because the auditory and visual sources are simultaneously active during the multimodal encoding task, their signals are correlated at the scalp level. This correlation is particularly problematic for auditory source localisation, because auditory sources are often weaker than visual sources due to their increased distance from the electrodes. This makes it harder for auditory sources to be cleanly separated from simultaneously active visual response. Therefore, a separate unimodal localiser task was run for auditory sources, but not for visual sources.

For the visual sources, we took advantage of the phase structure of the audiovisual ratings task: because auditory stimuli are presented at either 0° or 180° starting phase, the auditory responses are in opposite phases and partially cancel each other out, resulting in reduced 4 Hz power at the auditory source. The visual stimulus, by contrast, always begins at the same phase, so its response adds constructively across trials, resulting in strong 4 Hz power at the visual source and providing a reliable estimate of visual source location. The spatial filters were therefore estimated using Linearly Constrained Minimum Variance (LCMV; Van Veen et al., 1997) beamformers separately using data in which the contribution of the other source was minimised, as described below.

For the auditory sources, sensor-level data from the unimodal localiser task were first transformed to Scalp Current Density (SCD) using the finite-difference method, and the same transformation was applied to the leadfields. This additional step prior to LCMV beamforming facilitates the localisation of two correlated, spatially separated auditory sources (Murzin et al., 2011). For the visual source, LCMV beamforming was applied directly to the sensor-level data from the audiovisual ratings task, without prior SCD transformation.

For both conditions, source-level data were averaged across trials to obtain event-related potentials (ERPs) before time-frequency analysis. Time-frequency analysis was applied to each source ERP using Morlet wavelets (width = 7 cycles) at frequencies between 3 and 5 Hz in steps of 0.5 Hz, and at time points between −2 and 5 s post-stimulus onset in steps of 0.05 s. Evoked power at each virtual electrode was averaged over 0.75-2.75 s post-stimulus onset within ±0.5 Hz of the stimulation frequency (3.5-4.5 Hz) and normalised using the pre-stimulus baseline period (-1.75 to -0.25 s relative to stimulus onset). Normalised evoked power was grand-averaged across participants and interpolated onto the MNI template MRI.

Two out of three regions of interest (ROIs) were identified as the locations of maximum grand-average evoked power: the right auditory cortex (MNI: [50, −20, 10] mm; Heschl’s gyrus R) and the visual cortex (MNI: [10, −100, 20] mm; Fig 2A-B). The left auditory ROI could not be reliably identified in the present dataset, as the peak evoked power fell outside expected anatomical boundaries with a substantially lower value than the right hemisphere. The left auditory ROI coordinates were therefore taken from Wang et al. (2023), who used the same stimulation protocol and stimulus sets in the unimodal condition (MNI: [-50,-31,0] mm). The validity of these coordinates was confirmed in two ways: first, the group-average ERP at the selected voxel showed clear 4-Hz activity in the unimodal auditory data; second, during multimodal source reconstruction, the reconstructed time series at this location exhibited a clear 4-Hz oscillation consistent with rhythmic entrainment to the auditory stimulus, as verified through visual inspection of the source-level evoked activity at both the group level (Fig 2C-D) and for individual participants (Supplementary Figures S1-2).

**Fig 1.**
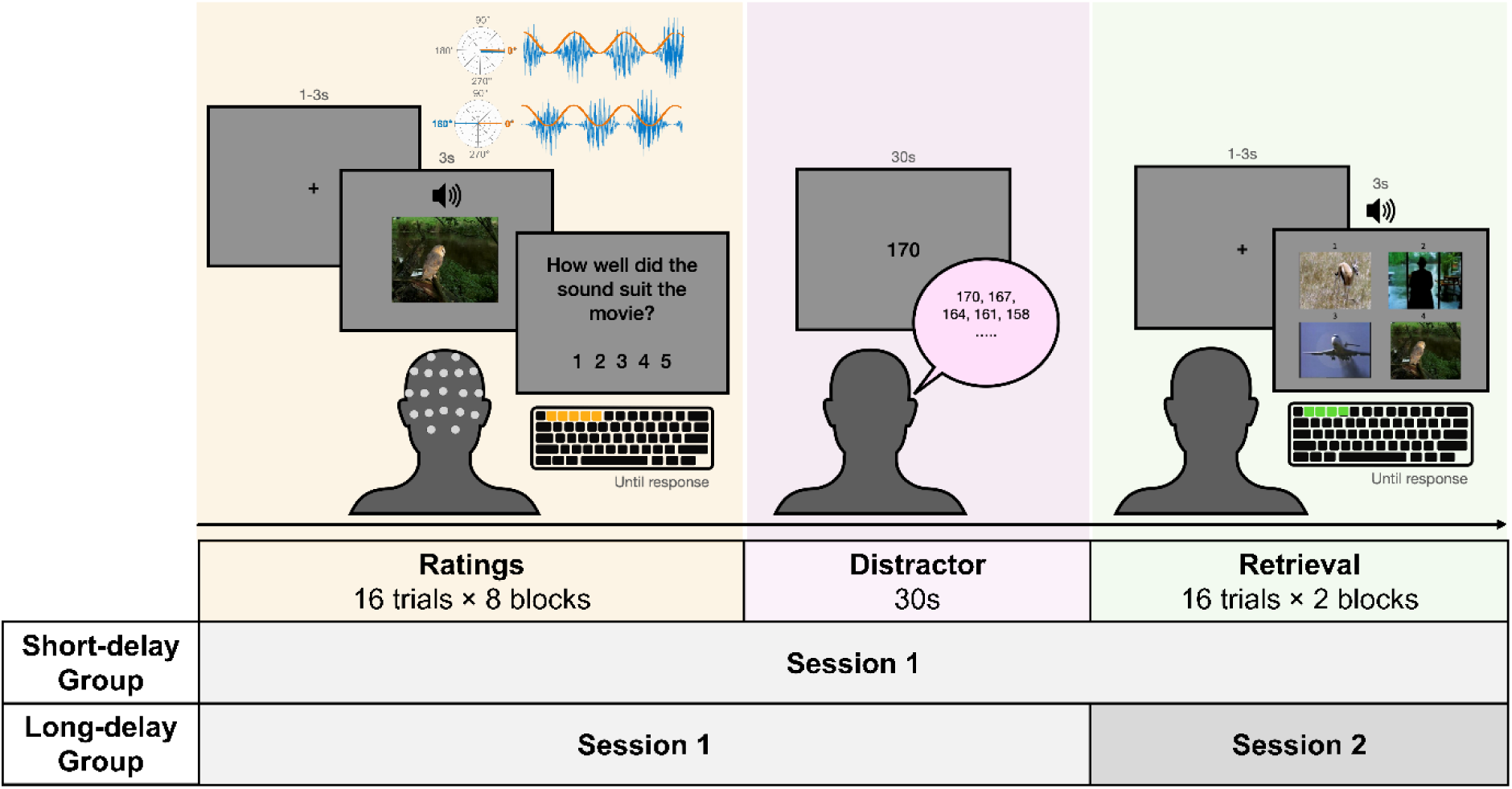
**Experimental paradigm and design**. During the encoding phase (ratings task), EEG was recorded while participants were presented with unrelated audio-video pairs and asked to rate how well the sound suited the video on a scale of 1 to 5. Both the auditory (blue) and visual stimuli (orange) were modulated at 4 Hz. The phase relationship between the two streams was manipulated across trials, such that they were presented either in synchrony (0° phase difference) or out of synchrony (180° phase offset). Each of the 32 pairs was presented 4 times across 8 blocks (16 trials per block). EEG was recorded during this phase. Following a brief distractor task (30 s of counting backward in steps of 3), participants completed a four alternative forced-choice memory test of the 32 pairs across 2 blocks (16 trials per block), in which participants selected the image previously paired with each sound. The short-delay group completed encoding, distractor, and retrieval within a single session, whereas the long-delay group completed encoding and distractor in Session 1 and retrieval in Session 2.

**Fig 2.**
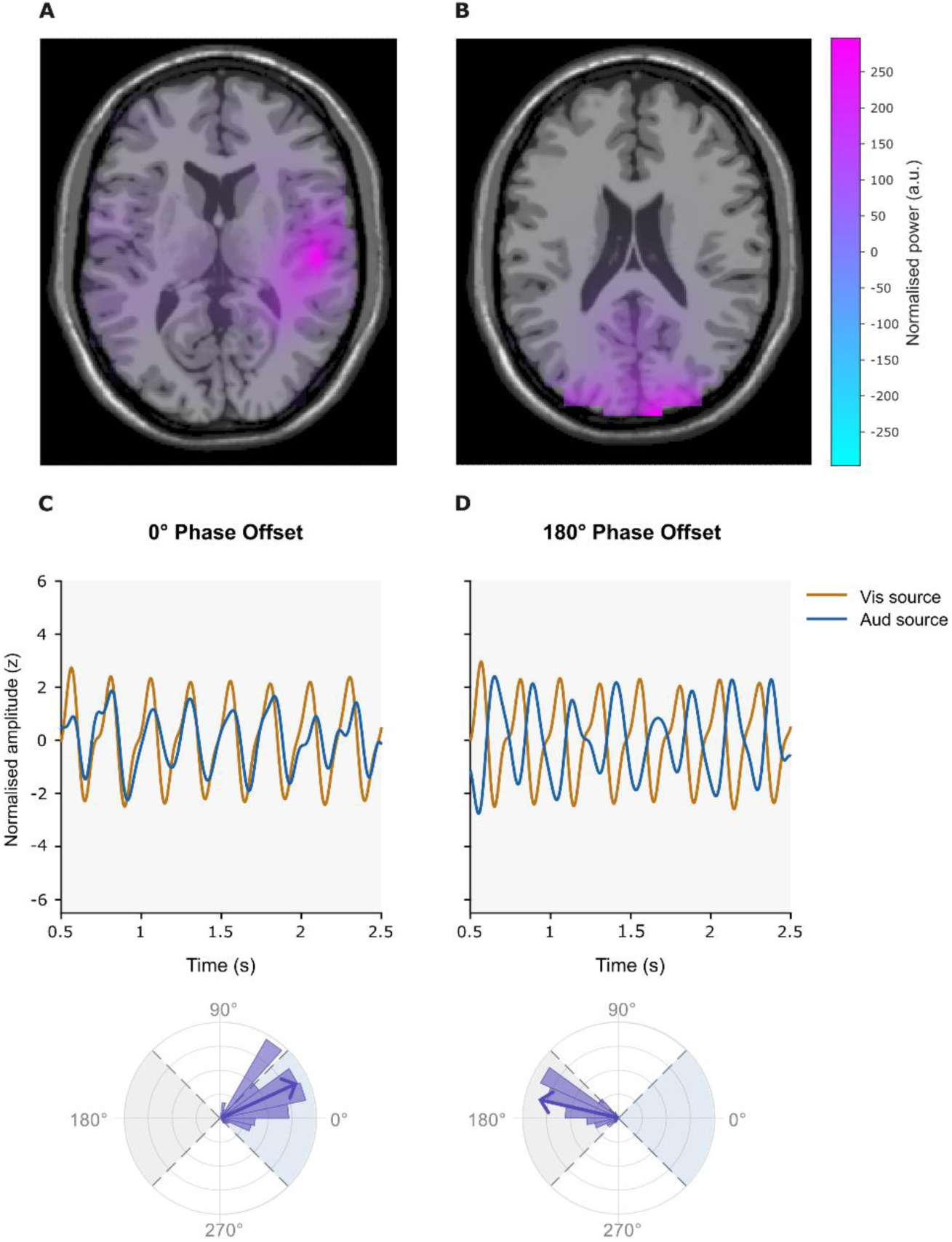
**Group-level source localisation of auditory and visual ROIs**. (A) Grand-average evoked power at 4 Hz (Morlet wavelets, 0.75–2.75 s post-stimulus onset, normalised to pre-stimulus baseline −1.75 to −0.25 s) projected onto the MNI template MRI, axial view. Peak voxel identified in right auditory cortex (Heschl’s gyrus R; MNI: 50, −20, 10 mm). Source reconstruction was performed on unimodal auditory data using LCMV beamforming on SCD-transformed data. (B) Grand-average evoked power at 4 Hz for the visual ROI. Peak voxel identified in occipital cortex (MNI: 10, −100, 20 mm). Source reconstruction was performed on multimodal RSS ratings data (not SCD-transformed) using LCMV beamforming. (C-D top) Grand average source-level ERPs (bandpass filtered 1.5–9 Hz, z-score normalised) in the analysis window (0.5-2.5 post-pair onset) at the auditory (light-blue) and visual (orange) ROIs for audio-video pairs assigned to the 0° phase bin (top-left) and the 180° phase bin (top-right). (C-D bottom) Rose histograms showing the distribution of instantaneous phase differences across all participants for the 0° phase bin (bottom-left) and the 180° phase bin (bottom-right). The resultant vector indicates the circular mean direction.

**Fig 3.**
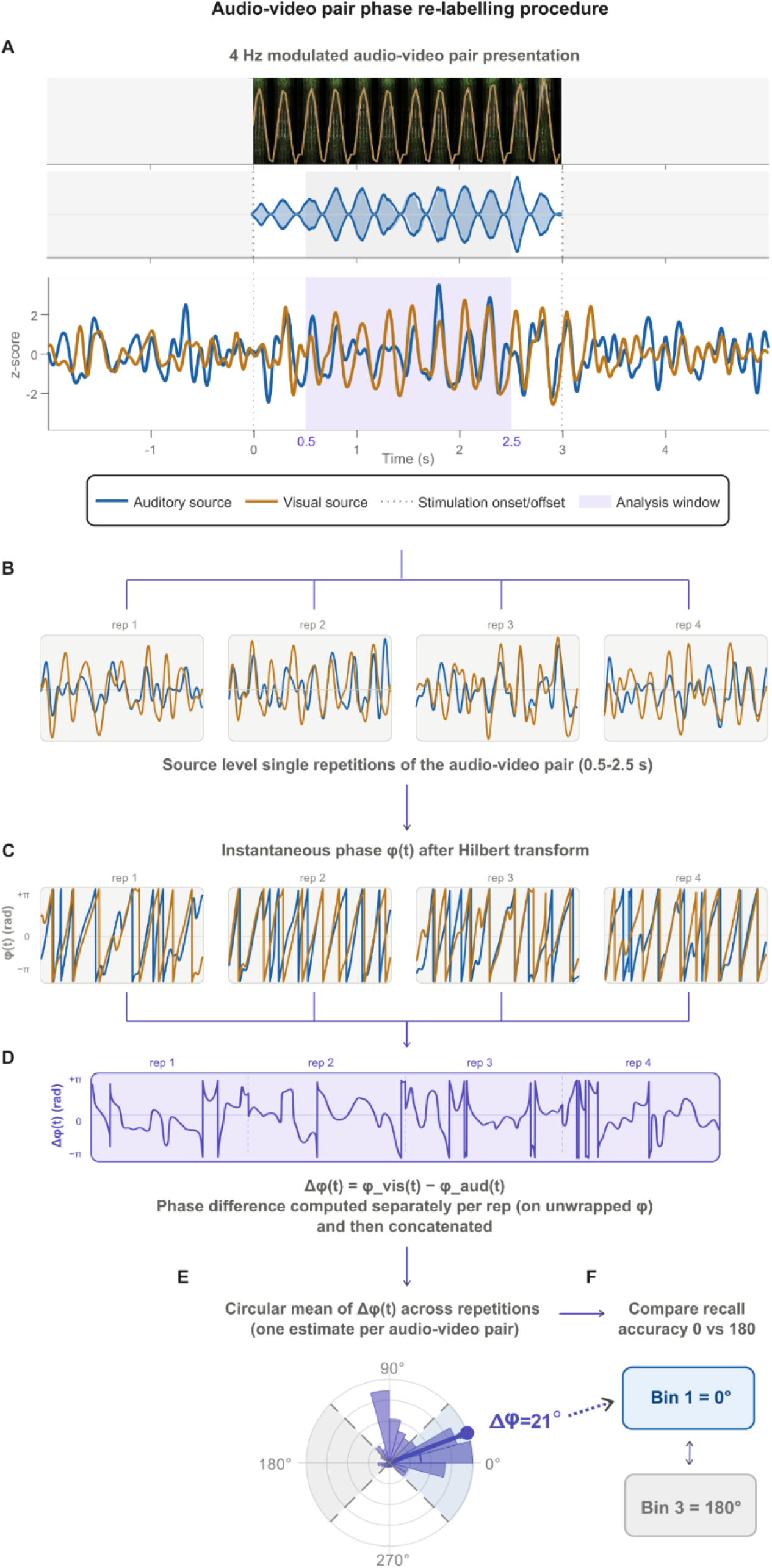
(A) Top: Example audiovisual pair consisting of a visual stimulus with a 4-Hz luminance modulation and an auditory stimulus with a 4-Hz amplitude modulation presented with a zero-phase offset. Bottom: Average source-level signals from the auditory and visual ROIs across the four presentations after bandpass filtering (1.5–9 Hz) and z-score normalization. The shaded region indicates the analysis window (0.5–2.5 s) on which the phase offset estimation between sources was performed. (B) Bandpass-filtered source-level signals from the auditory and visual ROIs after the multimodal source reconstruction for each of the four presentations within the analysis window. (C) Instantaneous phase of each source signal estimated using the Hilbert transform. (D) The instantaneous phase difference between the visual and auditory signals is computed separately for each presentation using the unwrapped phase estimates and subsequently concatenated across the four repetitions. (E) A circular mean phase difference was calculated across all concatenated time points to obtain a single phase estimate for each audiovisual pair. The rose histogram illustrates the distribution of instantaneous phase differences across all concatenated time points for the example audiovisual pair. The resultant vector (arrow) defines the pair’s circular mean phase difference, which is used for phase-bin assignment. Each pair is assigned to one of four phase bins centred at 0°, 90°, 180°, or 270° according to its circular phase difference. (F) Memory performance is then compared between audiovisual pairs assigned to opposite phase bins (0° versus 180°).

**Fig 4.**
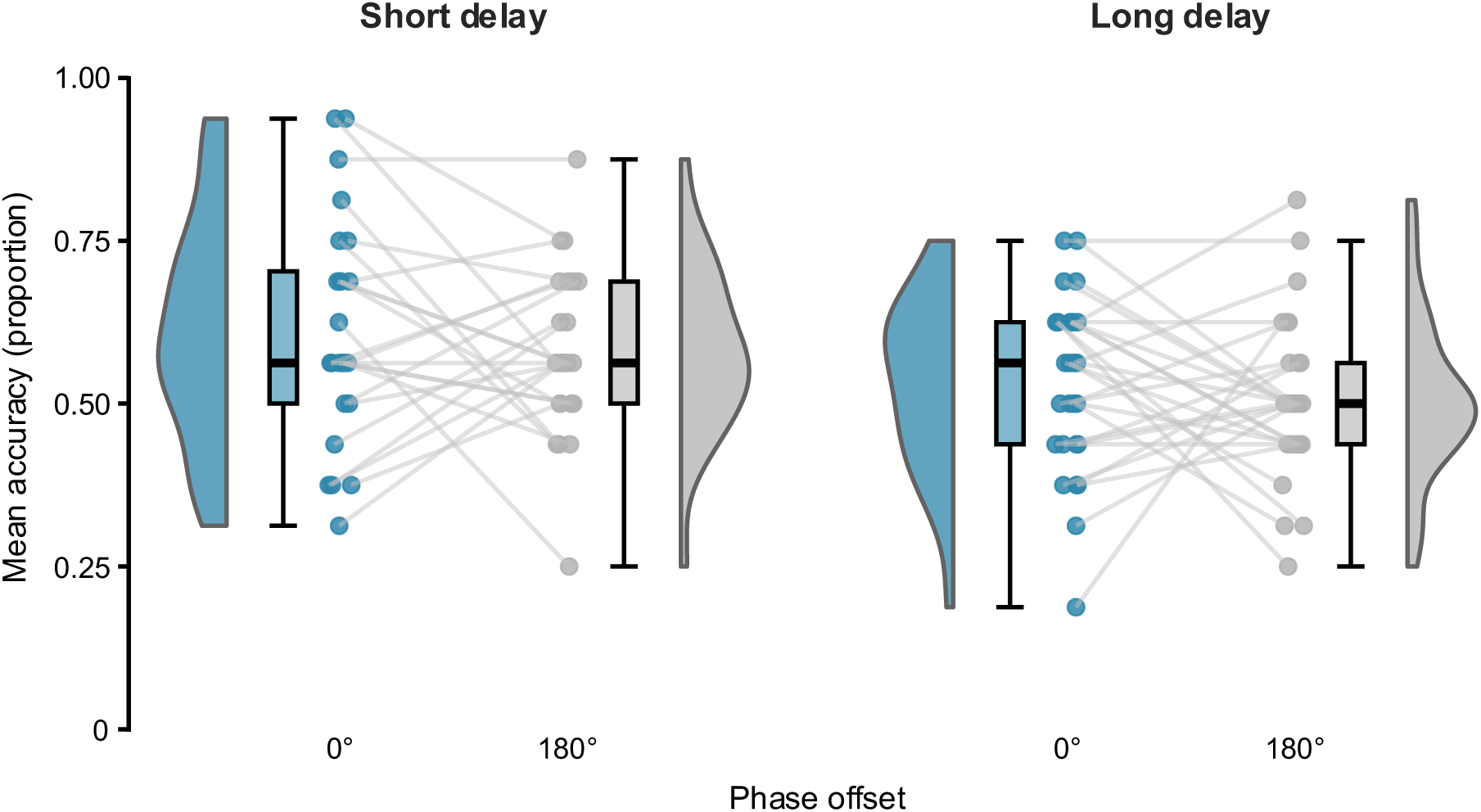
**Behavioural memory performance by phase offset and group**. Raincloud plots depicting the difference in accuracy between 0 and 180 phase offsets in the short-delay and long-delay groups. Coloured circles represent participants’ proportion of correctly remembered pairs. Boxplots show median and interquartile range.

**Fig 5.**
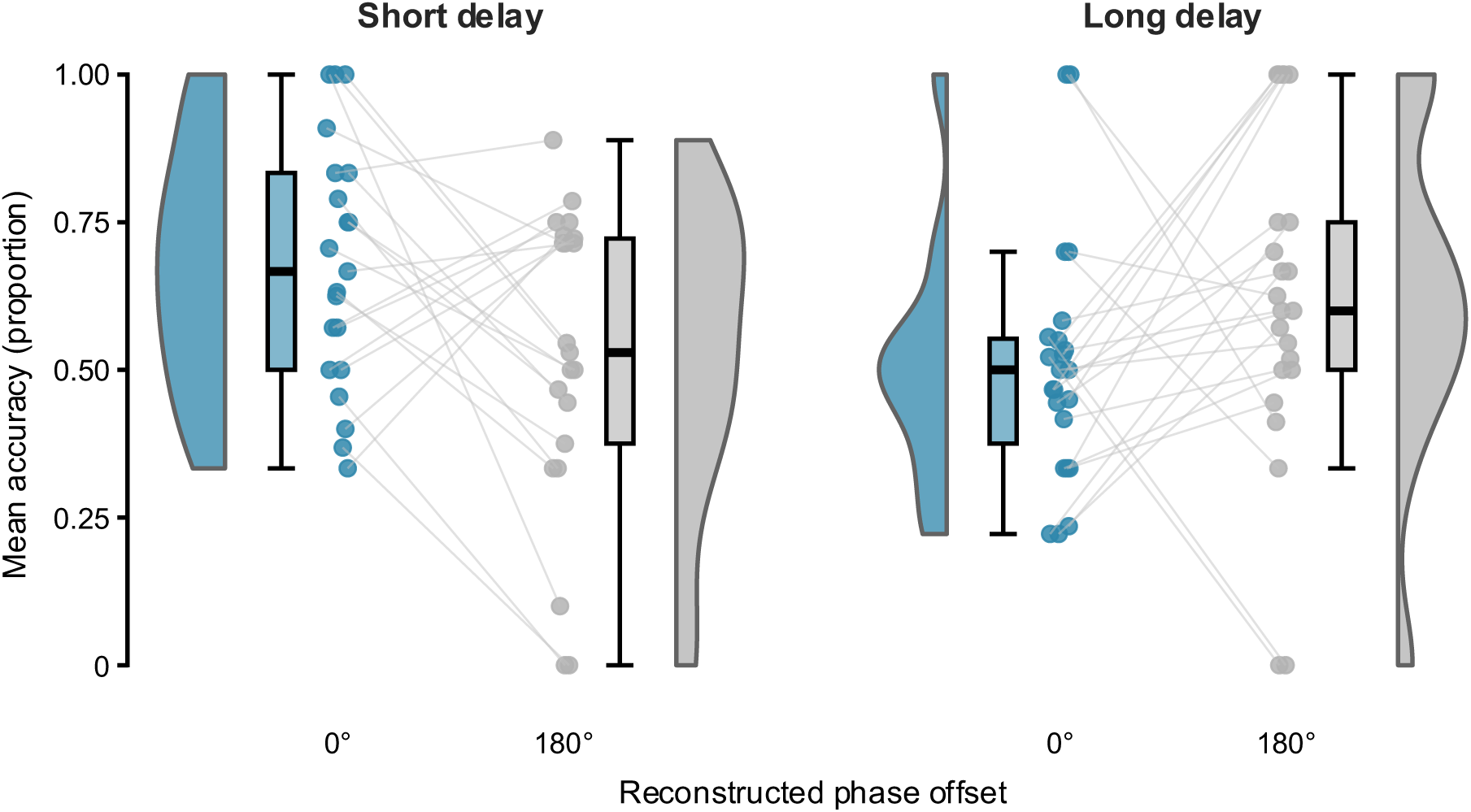
**Memory performance by reconstructed phase offset and group**. Raincloud plots depicting the difference in accuracy between 0 and 180 reconstructed phase bins in the two groups. Coloured circles represent participants’ proportion of correctly remembered pairs. Boxplots show median and interquartile range.

#### 2.5.3 Multimodal source localization

Source-level time series were reconstructed at the three identified ROIs using LCMV beamforming applied to the multimodal encoding (ratings task) data. For the two auditory ROIs, sensor-level data were first SCD-transformed, and separate sets of spatial filters were computed for the left and right hemispheres. The time series at the visual ROI were reconstructed directly with LCMV without prior SCD transformation. As LCMV beamforming results are inherently ambiguous regarding dipole orientations, the signs of the reconstructed time series were manually adjusted for each participant based on visual inspection of the spatial filter topography. The spatial filters were saved prior to sign flipping to ensure reproducibility. The time series from the left and right auditory ROIs were subsequently averaged to obtain a single auditory source signal per participant.

#### 2.5.4 Trial-level phase back-sorting

The source-level time series were bandpass filtered between 1.5 and 9 Hz and Hilbert-transformed to extract instantaneous phase angles. Since each audio-video pair was presented four times during encoding, the phase angle time points from all surviving repetitions after preprocessing of the same pair were concatenated before computing the circular mean, effectively increasing the number of time points available for phase estimation. The mean phase difference between the visual and auditory ROIs was then computed across data points within 0.5–2.5 s post-stimulus onset. Trials were sorted by mean phase difference from −π to π and assigned to one of four phase bins (0°, 90°, 180°, 270°) using fixed bin boundaries of ±45° around each phase of interest. This method ensures accurate trial assignment at the cost of unequal trial numbers across bins, which was addressed in the statistical analysis by using trial-level GLMMs. Memory accuracy was computed separately for each phase bin, and the 0° and 180° bins were used in all subsequent statistical analyses.

### 2.6 Statistical analysis

All statistical analyses were conducted in R (version 4.5.2; R Core Team, 2025), and statistical significance was defined as p < .05.

To examine the effects of stimulus phase offset (0° vs. 180°) and group (short vs. long delay) on memory accuracy, mixed ANOVAs were conducted with phase offset as a within-subjects factor and group as a between-subjects factor, using Type III sums of squares. Assumptions of normality and homogeneity of variance were assessed through visual inspection of Q–Q plots and Levene’s test, respectively. Effect sizes for ANOVA effects are reported as partial eta squared (η²p). For the behavioural analyses, descriptive within-group differences between the 0° and 180° phase-offset conditions are accompanied by paired Cohen’s d_z_ values with 95% confidence intervals (CI).

The same analytical approach was applied to the source-reconstructed phase-offset data. Participants who did not contribute observations to both reconstructed phase bins were excluded from these analyses. Significant interactions were followed by post hoc pairwise comparisons of estimated marginal means. Resulting p-values were corrected for multiple comparisons using the false discovery rate (FDR; Benjamini & Hochberg, 1995) procedure, and effect size reported as Cohen’s d with 95% confidence intervals. Within-group phase comparisons (0° vs. 180°) in the short-delay group were tested one-tailed based on the directional hypothesis that memory accuracy would be higher at 0° than 180° reconstructed phase, consistent with previous studies. In contrast, comparisons in the long-delay group were evaluated using two-tailed tests because two competing hypotheses were considered regarding the effect of overnight consolidation (i.e., that the synchrony-related memory advantage could either persist or be attenuated).

A generalised linear mixed model (GLMM) was fitted to the binary trial-level data (hit vs miss) using the lme4 package (Bates et al., 2015). The model included reconstructed phase offset and group as fixed effects, with predictor variables contrast-coded and mean-centred prior to analysis. Phase offset was coded such that 0° = positive and 180° = negative, and group was coded such that short delay = positive and long delay = negative. Random effects structure was kept maximal following Barr et al. (2013), including random intercepts and slopes for both participants and stimuli (movies and sounds); all models, including progressively simplified random effects structures, were singular but converged successfully. We therefore retained the maximal model as suggested by Barr et al., 2013 and Matuschek et al., 2017. Model fitting used the optimx optimizer with the L-BFGS-B method. Fixed effects are reported as unstandardised logistic regression coefficients (β) with standard errors, z-statistics, p-values, and 95% Wald confidence intervals.

To assess whether participants could consciously discriminate between synchronous and asynchronous audiovisual pairs in the synchronisation task, sensitivity (d′) was computed for each participant from their synchronisation task responses, using hits (correct identification of synchronous pairs) and false alarms (incorrect identification of asynchronous pairs as synchronous). A one-sample t-test was used to assess whether mean d′ differed significantly from zero across all participants (n = 63). To examine whether individual differences in conscious synchrony detection could account for the observed phase offset effects on memory, d′ was subsequently included as a continuous covariate in the mixed ANOVAs for both the behavioural (n = 52) and source-reconstructed (n = 44) datasets. Where a significant d′ × phase offset interaction was observed, simple contrasts were computed at ±1 SD of d′ around the mean using the emmeans package. Additionally, d′ was added as a fixed effect to the GLMM to assess its influence at the trial level and to examine whether its inclusion improved model fit, assessed via likelihood ratio test comparing models with and without d′.

Mixed ANOVAs were implemented using the afex package (Singmann et al., 2025), estimated marginal means and post hoc comparisons using emmeans (Lenth & Piaskowski, 2026), and effect sizes using effectsize (Ben-Shachar et al., 2020).

## 3 Results

### 3.1 Behavioural memory performance

To examine whether the phase offset between audio and video stimuli (0° vs 180°) influenced memory, and whether this differed between groups, we conducted a mixed ANOVA with phase offset (0° vs 180°) as a within-subjects factor and group (short vs long delay) as a between-subjects factor.

The ANOVA revealed a significant main effect of group (F(1, 50) = 6.75, p = .012, η²p = .12), with the short-delay group showing higher accuracy (M = 0.595, SE = 0.022) than the long-delay group (M = 0.519, SE = 0.020). There was no significant main effect of phase offset (F(1, 50) = 1.21, p = .276, η²p = .02) and no significant phase offset x group interaction (F(1, 50) = 0.03, p = .861, η²p = < .001). Although not statistically significant, the average memory accuracy was higher for 0° than 180° phase offset in both groups (short-delay: M₀ = 0.612, SD = 0.176; M₁₈₀ = 0.578, SD = 0.131, Cohen’s d_z_ = 0.17, 95% CI [-0.23, 0.57]; long-delay: M₀ = 0.531, SD = 0.134; M₁₈₀ = 0.507, SD = 0.125, Cohen’s d_z_ = 0.13, 95% CI [-0.24, 0.51]).

### 3.2 Effects of reconstructed theta phase offset and group on memory accuracy

To test whether memory accuracy differed as a function of the reconstructed phase offset in the sensory cortices at encoding (0° vs 180°), and whether this differed between groups, we conducted a mixed ANOVA with reconstructed phase (0° vs 180°) as a within-subjects factor and group (short vs long delay) as a between-subjects factor. Eight participants were excluded from this analysis due to missing data in one of the two phase bins (short delay: n = 3; long delay: n = 5), resulting in a final sample of 44 participants (short delay: n = 21; long delay: n = 23). The ANOVA revealed a significant group x reconstructed phase interaction (F(1, 42) = 6.08, p = .018, η²p = .126), with no significant main effect of group (F(1, 42) = 0.60, p = .443, η²p = .014) or reconstructed phase (F(1, 42) = 0.17, p = .686, η²p = .004).

To unpack the interaction, pairwise comparisons were conducted with FDR correction. Within the short-delay group, memory accuracy was significantly higher at 0° (M = 0.676, SE = 0.045) than 180° reconstructed phase offset (M = 0.519, SE = 0.059; t(42) = 1.987, p = .027, one-tailed, d = 0.47, 95% CI for Cohen’s d [0.01, 0.92]). Within the long-delay group, there was no significant difference between 0° (M = 0.504, SE = 0.043) and 180° reconstructed phase (M = 0.617, SE = 0.056; t(42) = -1.490, p = .144, two-tailed, d = -0.29, 95% CI for Cohen’s d [-0.71, 0.13]).

At 0° reconstructed phase, the short-delay group showed significantly higher memory accuracy than the long-delay group (mean difference = 0.172; t(42) = 2.76, p = .009). In contrast, the groups did not differ significantly in the 180° phase condition (mean difference = -0.098, t(42) = -1.20, p = .237).

Given that the number of trials per reconstructed phase bin varied considerably across participants, with some participants contributing as few as 1 trial to a bin (short-delay group, 0° bin: M = 10.05, SD = 7.13, range = 1–26; 180° bin: M = 9.87, SD = 6.70, range = 1–25; long-delay group, 0° bin: M = 13.77, SD = 7.82, range = 1–29; 180° bin: M = 7.72, SD = 7.30, range = 1–27), we complemented the ANOVA with a GLMM on trial-level binary memory outcomes (hit vs miss). This approach avoids aggregating data into potentially unreliable bin-level accuracy scores and accounts for both participant- and stimulus-level variance through random effects. At the trial level, the GLMM showed an interaction estimate in the same direction as the ANOVA; however, the interaction only approached significance (β = -0.694, SE = 0.357, z = -1.943, p = .052, 95% CI [-1.395, 0.006]), indicating a trend toward differential effects of reconstructed phase across groups. No significant main effect of reconstructed phase was observed (β = -0.184, SE = 0.190, z = -0.971, p = .332), consistent with the opposing directions of the phase effect in the two groups. Finally, a significant main effect of group was observed (β = -0.365, SE = 0.157, z = -2.325, p = .020, 95% CI [-0.673, -0.057]), indicating higher overall memory performance in the short-delay group and consistent with the main effect of group observed in the behavioural analysis. To further examine the interaction, separate GLMMs were fitted within each group. In the short-delay group, the reconstructed phase effect was in the expected direction (higher memory accuracy at 0° than 180°), but was not statistically significant (β = 0.245, SE = 0.298, z = 0.822, p = .206, one-tailed, 95% CI [−0.339, 0.829]). In the long-delay group, reconstructed phase significantly predicted memory accuracy (β = −0.512, SE = 0.233, z = −2.203, p = .028, 95% CI [−0.968, −0.057]), indicating higher memory accuracy at 180° than 0°.

Because subjective ratings of the stimuli may influence subsequent memory performance, we conducted an additional generalized linear mixed-effects analysis including the subjective rating as a covariate. This model tested whether the observed effects of reconstructed phase and group persisted after accounting for participants’ ratings of the stimuli. Consistent with previous findings (Wang et al., 2026), higher subjective ratings predicted an increased likelihood of remembering (β = 0.26, SE = 0.07, z = 3.62, p < .001, 95% CI [0.12, 0.40]). Importantly, controlling for ratings did not meaningfully alter the estimated Reconstructed Phase × Group interaction (β = -0.67, SE = 0.35, z = -1.89, p = .059, 95% CI [-1.36, 0.03]), which remained very similar in magnitude to that observed in the model excluding ratings (β = -0.69). Including subjective ratings did not materially change the magnitude of the estimated interaction, suggesting that differences in subjective ratings are unlikely to fully explain the observed phase-dependent group differences.

### 3.3 Audiovisual synchrony discrimination effects on memory performance

To verify that participants could not consciously discriminate between the synchronous and asynchronous conditions, the synchronisation task was analysed across all participants (n = 63).. Sensitivity (d’) was computed for each participant based on hits (correctly identifying a synchronous pair as synchronous) and false alarms (incorrectly identifying an asynchronous pair as synchronous). Mean d’ did not differ significantly from zero (M = 0.13, SD = 0.61, 95% CI for the mean d′ [−0.03, 0.28], t(62) = 1.657, p = .103, Cohen’s d = 0.21, 95% CI [−0.04, 0.46]), suggesting that participants were unable to reliably detect the phase manipulation.

Although mean d′ did not differ significantly from zero, the result approached significance. Therefore, to examine whether individual differences in explicit synchrony discrimination influenced memory performance, d′ was included as a continuous covariate in the behavioural mixed ANOVA (n = 52, above guessing threshold). Neither the main effect of d′ (F(1, 49) = 1.00, p = .322, η²p = .020) nor its interaction with phase offset (F(1, 49) = 0.10, p = .749, η²p = .002) was significant. The main effect of group remained significant (F(1, 49) = 6.90, p = .011, η²p = .123), indicating that the behavioural results were not explained by participants’ conscious sensitivity to the level of synchrony of the stimuli.

We then repeated this control analysis for the source-reconstructed dataset (n = 44). Including d′ as a covariate revealed a significant main effect of d′ (F(1,41) = 6.49, p = .015, η²p = .137) and a significant d′ × reconstructed phase interaction (F(1,41) = 5.65, p = .022, η²p = .121). Higher d′ was associated with higher overall memory accuracy (r = .35, p = .018), an association driven primarily by memory in the 180° reconstructed phase (r = .46, p = .002), with no association for the 0° reconstructed phase (r = −.06, p = .715). Simple contrasts revealed that the expected memory advantage for 0° over 180° pairs was significant only at low d′ (-1 SD; estimate = 0.185, p = .037) and absent at mean d′ (estimate = 0.022, p = .677). At high d′ (+1 SD), the pattern numerically reversed, with 180° pairs showing higher accuracy than 0° pairs, although this did not reach significance (estimate = −0.142, p = .108). However, this d′ × phase interaction was not replicated in the trial-level GLMM (β = −0.289, z = −0.959, p = .338), and including d′ in the GLMM did not improve model fit (χ²(1) = 0.651, p = .420). Importantly, the group × reconstructed phase interaction remained stable before and after controlling for d′ across all models (ANCOVA: F(1,41) = 5.62, p = .023, ηp² = .121; GLMM without d′: β = −0.694, p = .052; GLMM with d′: β = −0.712, p = .048), confirming that the interaction was not attributable to individual differences in conscious awareness of the degree of audiovisual synchrony.

## 4 Discussion

The present study investigated whether the Theta-Induced Memory Effect (TIME), i.e. the enhancement of associative memory accuracy observed when visual and auditory stimuli to be encoded are rhythmically modulated at theta frequency and presented in-phase compared to out-of-phase, persists following overnight consolidation. At the behavioural level, no average-level differences were observed between in-phase and out-of-phase stimulation. However, source-level back-sorting based on reconstructed phase differences replicated the TIME effect in the short-delay group. In contrast, the long-delay group exhibited a reversed pattern, suggesting that the relationship between multisensory phase synchrony at encoding and subsequent associative memory changes over time.

The absence of a behavioural effect at the average level is consistent with previous studies (Serin et al., 2024; Wang et al., 2026). As demonstrated by Wang et al. (2026), the experimentally imposed phase offset does not necessarily correspond to the phase relationship achieved at the sensory cortices on individual trials, owing to variability in neural entrainment. Such variability may arise from individual differences in sensory transduction delays and fluctuations in ongoing brain state and, in the present study, may have been further amplified by the repeated presentation of each audio-visual pair. Specifically, each pair was presented four times across eight blocks, potentially introducing additional variability in entrainment efficacy across repetitions. Consequently, averaging behavioural performance across trials may obscure phase-dependent memory effects that only emerge when the actual phase relationship is reconstructed at the source level. The trial-level back-sorting procedure addresses this issue by reclassifying trials according to the reconstructed phase difference between auditory and visual cortices. In the present study, this phase difference was estimated by concatenating phase signals across the four repetitions and calculating their circular mean, which may explain why the TIME effect was detectable after back-sorting but not before.

Importantly, back-sorting trials based on source reconstructed phase differences produced the expected phase-memory relationship in the short-delay group, reinforcing previous evidence that memory formation depends on the phase relationship achieved at sensory cortices rather than the phase offset imposed experimentally (Wang et al., 2018, 2026).In contrast, the long-delay group showed a tendency towards a reversed phase-memory relationship. Although the statistical significance of the within-group phase effects differed between the participant-level ANOVA and the trial-level GLMM, both analyses converged on the same qualitative interaction: memory performance was higher following the 0° than the 180° reconstructed phase in the short-delay group, whereas the opposite pattern was observed following the overnight delay. The differing significance of the simple effects likely reflects the limited statistical power of the present study, together with the substantial imbalance in the number of reconstructed trials contributing to each phase bin. Whereas the ANOVA assigns equal weight to each participant, the GLMM operates at the trial level and therefore weights participants according to the number of reconstructed trials contributing to each phase bin.

Overall memory performance was significantly lower in the long-delay group than in the short-delay group, consistent with the well-established forgetting of incidentally encoded and unrehearsed information over time (Ebbinghaus, 1885; Murre & Dros, 2015). More interestingly, however, the present findings suggest that the relationship between encoding-related synchrony and subsequent memory may itself change across consolidation.

One possible interpretation is that theta synchrony primarily facilitates the initial formation of associative representations, whereas the long-term maintenance of those representations depends on subsequent consolidation processes. Under this account, synchrony-related benefits established during encoding do not necessarily translate into a durable memory advantage, but may instead be modified as hippocampus-dependent memories undergo offline consolidation. Contemporary accounts propose that newly acquired memories undergo transformation through repeated reactivation and integration with existing knowledge structures (Dudai et al., 2015; Klinzing et al., 2019). Consistent with this view, replay during rest and sleep is increasingly recognised as a selective process that preferentially strengthens only a subset of newly acquired memories, rather than uniformly consolidating all encoded experiences. Which memories are preferentially replayed remains an active area of investigation. Some accounts suggest that replay prioritises memories that are particularly relevant for future behaviour (Ólafsdóttir et al., 2018), and more recent work suggests that replay probability may be governed by dynamic episode-specific priority traces that evolve over time, rather than simply reflecting the reward value of an experience (Tirole et al., 2026, preprint). Given that encoding in the present study was incidental, participants had no explicit expectation that the associations would later be tested, potentially reducing the behavioural relevance of the encoded material. Consequently, synchrony-dependent memory traces may have weakened or been reorganised over time and may not have been preferentially reactivated during consolidation. In addition to replay-based accounts, sleep has also been proposed to renormalise synaptic strength through global downscaling (Tononi & Cirelli, 2006, 2014). If the synchrony-related memory advantage reflects greater synaptic potentiation during encoding, such renormalisation could have further reduced the difference between 0 and 180 phase degree offsets trials established during encoding. Also, the phase reversal in the long-delay group could be explained by some behavioural studies that have reported preferential sleep-related benefits for relatively weaker memories under particular experimental conditions (Petzka et al., 2021), although this effect appears to depend on factors such as retrieval demands and initial memory strength and may not generalise across paradigms. Because the present study did not directly manipulate or measure these factors, it remains unclear whether such mechanisms contributed to the current findings. Several limitations should be acknowledged. First, source reconstruction reduced the sample size from 52 to 44 participants, limiting statistical power. In addition, the back-sorting procedure distributed a relatively small number of trials across four phase bins, resulting in unequal trial counts and, in some cases, extreme accuracy estimates. These factors likely contributed to the variability in the statistical significance of the ANOVA and GLMM despite their convergence in the overall qualitative pattern. More broadly, the study was designed in 2023 and data collection was completed in 2024, before several important replication attempts had been published. Subsequent studies have revealed considerable variability in the magnitude and reliability of the TIME effect (Serin et al., 2024; Simeonov & Das, 2025; Wang et al., 2026), suggesting that larger samples may be required to obtain stable estimates of synchrony-dependent memory effects. Finally, sleep was assessed only by self-report, and no sleep physiology or replay-related neural activity was measured, precluding direct conclusions regarding the mechanisms underlying the observed changes across consolidation. Future studies should test larger samples and include objective measures of sleep to better characterise the temporal dynamics of the TIME effect. Examining multiple retrieval intervals may help determine whether the synchrony-related memory advantage decays gradually or undergoes a qualitative reorganisation across consolidation. In addition, combining EEG source reconstruction with sleep measures may provide valuable insights into whether sleep-related processes contribute to the persistence or transformation of synchrony-dependent memories. Finally, the use of individualised stimulation frequencies may help reduce variability in entrainment and further improve the reliability of phase-based memory effects.

In conclusion, the present study extends previous work by showing that the relationship between reconstructed multisensory theta phase at encoding and subsequent associative memory depends on when retrieval is tested. Whereas the expected phase-memory relationship was observed immediately after encoding, the opposite qualitative pattern emerged following an overnight delay. Together, these findings suggest that theta synchrony may primarily facilitate the initial formation of hippocampus-dependent associative memories, whereas the long-term expression of this encoding advantage depends on subsequent consolidation processes. Understanding how oscillatory synchronisation interacts with consolidation processes may therefore represent an important direction for future research.

## Data and Code Availability

The data analysed in this study are publicly available from OpenNeuro under the following link: https://doi.org/10.18112/openneuro.ds008095.v1.0.0. All analysis scripts used to preprocess the data and perform the analyses are available on GitHub at: https://github.com/NoT-CoOLab/longevity_TIME_EEG. The source reconstruction pipeline used to back-sort the trials is publicly available and can be found following this link: https://github.com/NoT-CoOLab/audiovisual-RSS-pipeline. Any further information required is available upon request.

## Author Contributions

Conceptualization, S.H., K.L.S., D.W., and E.M.; investigation, E. M.; formal analysis, and writing – original draft, E.M.; writing – review & editing, E.M., D.W., K.L.S., and S.H.; funding acquisition, S.H.; supervision, S.H. and K.L.S.

## Acknowledgements

The authors would like to thank Adela Maria Ostaf, Merwin Antonysiluvai, Ruiyao Tang, and Hana Hailu for their contribution to participant recruitment and data collection as part of their Master’s dissertation training and related research projects.

## Supplementary Materials

**Supplementary Figure S1.**
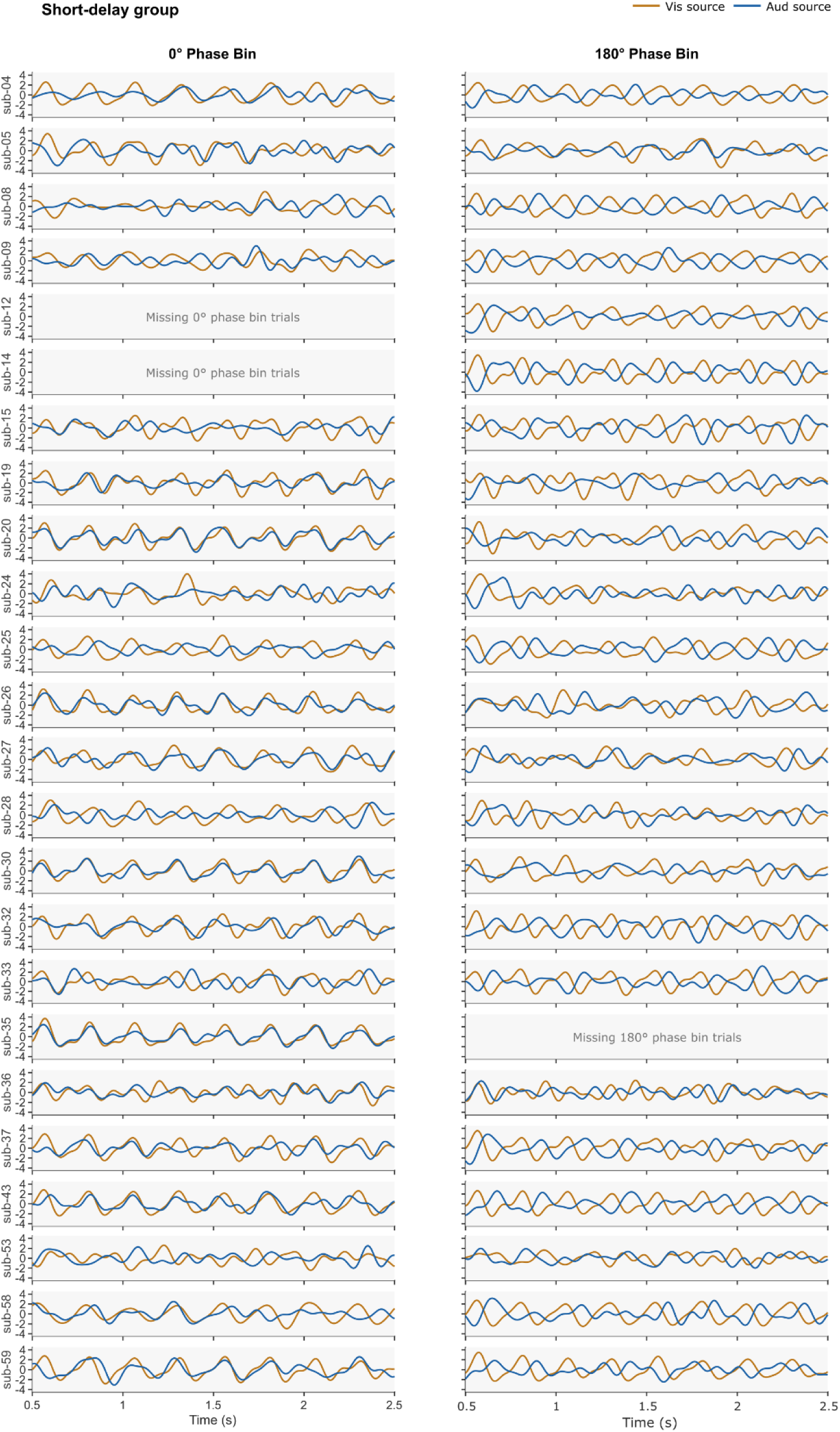
Individual participant source-level signals by phase bin in the short-delay group (n = 24). Average source-level ERP (bandpass filtered 1.5–9 Hz, z-score normalised) at the auditory and visual ROIs for each participant, shown separately for audio-video pairs assigned to the 0° phase bin (left column) and the 180° phase bin (right column). The auditory source signal represents the average of the left and right auditory ROIs source-reconstructed signals. Blank panels indicate that no trials were available for the corresponding reconstructed phase bin, resulting in exclusion of that participant from analyses requiring both phase bins. In the short-delay group, 3 participants had missing trials (0° phase bin: n = 2; 180° phase bin: n = 1).

**Supplementary Figure S2.**
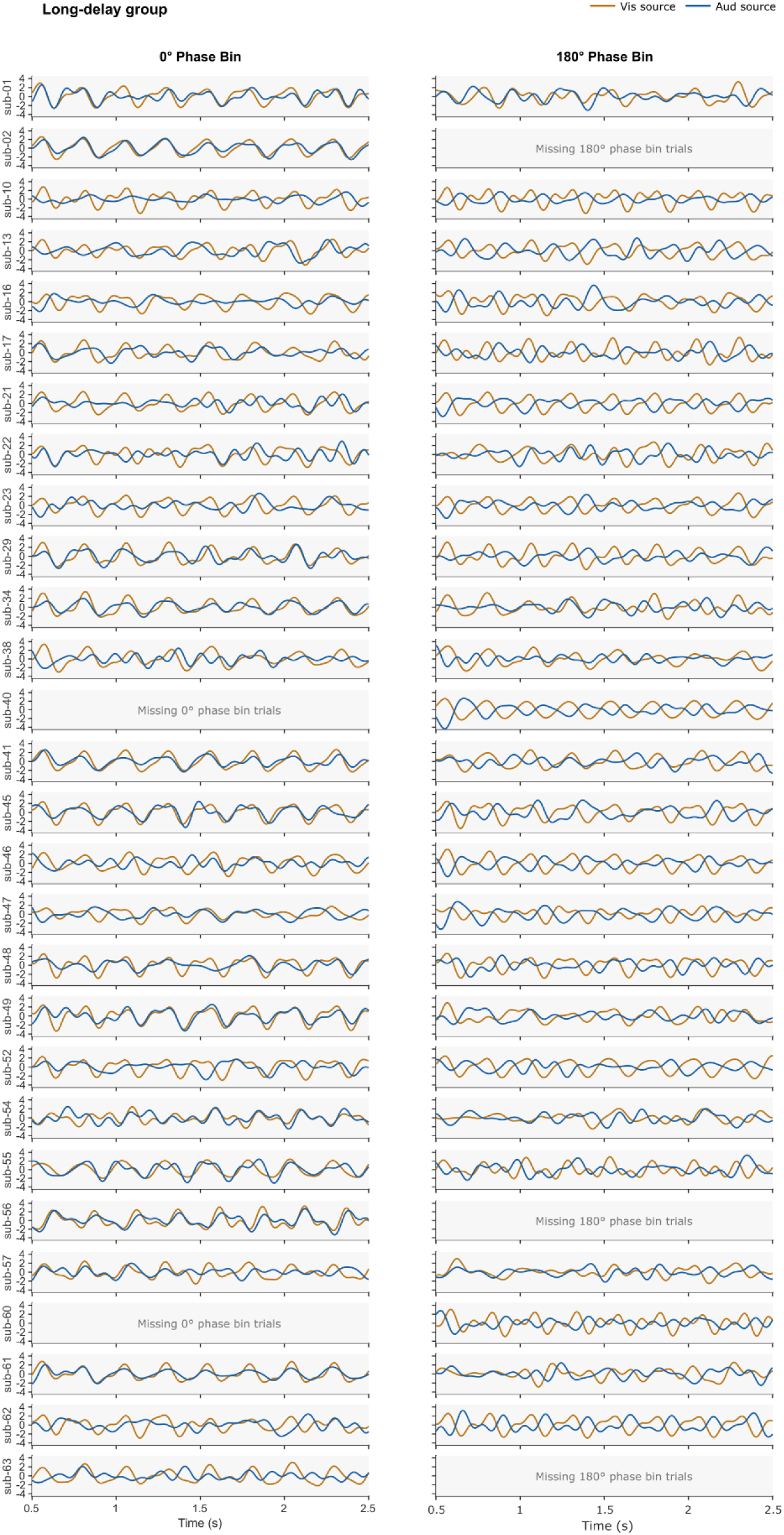
Individual participant source-level signals by phase bin in the long-delay group (n = 28). Same as Supplementary Figure S1, for participants in the long-delay group. Five participants had no trials reconstructed in one of the two phase bins (0° phase bin: n = 2; 180° phase bin: n = 3).

## References

1. Bates, D., Mächler, M., Bolker, B., & Walker, S. (2015). Fitting Linear Mixed-Effects Models Using lme4. *Journal of Statistical Software*, 67, 1–48. 10.18637/jss.v067.i01

2. Ben-Shachar, M. S., Lüdecke, D., & Makowski, D. (2020). effectsize: Estimation of Effect Size Indices and Standardized Parameters. Journal of Open Source Software, 5(56), 2815. 10.21105/joss.02815

3. Biau, E., Wang, D., Park, H., Jensen, O., & Hanslmayr, S. (2025). Neocortical and Hippocampal Theta Oscillations Track Audiovisual Integration and Replay of Speech Memories. Journal of Neuroscience, 45(21). 10.1523/JNEUROSCI.1797-24.2025

4. Bliss, T. V. P., & Lømo, T. (1973). Long-lasting potentiation of synaptic transmission in the dentate area of the anaesthetized rabbit following stimulation of the perforant path. The Journal of Physiology, 232(2), 331–356. 10.1113/jphysiol.1973.sp010273

5. Brainard, D. H. (1997). The Psychophysics Toolbox. Spatial Vision, 10(4), 433–436.

6. Clouter, A., Shapiro, K. L., & Hanslmayr, S. (2017). Theta Phase Synchronization Is the Glue that Binds Human Associative Memory. Current Biology, 27(20), 3143–3148.e6. 10.1016/j.cub.2017.09.001

7. Diekelmann, S., & Born, J. (2010). The memory function of sleep. Nature Reviews Neuroscience, 11(2), 114–126. 10.1038/nrn2762

8. Dudai, Y., Karni, A., & Born, J. (2015). The Consolidation and Transformation of Memory. Neuron, 88(1), 20–32. 10.1016/j.neuron.2015.09.004

9. Ebbinghaus, H. (1885). *Über das Gedächtnis: Untersuchungen zur experimentellen Psychologie*. Duncker & Humblot.

10. Hainke, L., Neumaier, V., Marcantoni, E., Wang, D., Capstick, K., Spitschan, M., Dowsett, J., & Hanlsmayr, S. (2026). *Audiovisual stimulation using wearable shutter glasses robustly evokes 40 Hz neuronal activity but does not modulate associative memory* (p. 2026.07.09.737418). bioRxiv. 10.64898/2026.07.09.737418

11. Hölscher, C., Anwyl, R., & Rowan, M. J. (1997). Stimulation on the Positive Phase of Hippocampal Theta Rhythm Induces Long-Term Potentiation That Can Be Depotentiated by Stimulation on the Negative Phase in Area CA1 In Vivo. The Journal of Neuroscience, 17(16), 6470–6477. 10.1523/JNEUROSCI.17-16-06470.1997

12. Huerta, P. T., & Lisman, J. E. (1995). Bidirectional synaptic plasticity induced by a single burst during cholinergic theta oscillation in CA1 in vitro. Neuron, 15(5), 1053–1063. 10.1016/0896-6273(95)90094-2

13. Hyman, J. M., Wyble, B. P., Goyal, V., Rossi, C. A., & Hasselmo, M. E. (2003). Stimulation in hippocampal region CA1 in behaving rats yields long-term potentiation when delivered to the peak of theta and long-term depression when delivered to the trough. The Journal of Neuroscience: The Official Journal of the Society for Neuroscience, 23(37), 11725–11731. 10.1523/JNEUROSCI.23-37-11725.2003

14. Kleiner, M., Brainard, D. H., Pelli, D., Ingling, A., Murray, R., & Broussard, C. (2007). What’s new in Psychtoolbox-3. Perception, *36*, 1–16. 10.1068/v070821

15. Klinzing, J. G., Niethard, N., & Born, J. (2019). Mechanisms of systems memory consolidation during sleep. Nature Neuroscience, 22(10), 1598–1610. 10.1038/s41593-019-0467-3

16. Lenth, R. V., & Piaskowski, J. (2026). emmeans: Estimated Marginal Means, aka Least-Squares Means. https://rvlenth.github.io/emmeans/

17. Marcantoni, E., Daube, C., Wang, D., Cao, C., Su, B., Zhan, S., Ince, R. A. A., Palva, S., Bush, D., & Hanslmayr, S. (2026). Non-invasive tracking of hippocampal theta oscillations. bioRxiv. 10.64898/2026.01.13.699218

18. Moscovitch, M. (2008). The hippocampus as a ‘stupid,’ domain-specific module: Implications for theories of recent and remote memory, and of imagination. Canadian Journal of Experimental Psychology = Revue Canadienne De Psychologie Experimentale, 62(1), 62–79. 10.1037/1196-1961.62.1.62

19. Murre, J. M. J., & Dros, J. (2015). Replication and Analysis of Ebbinghaus’ Forgetting Curve. PLoS ONE, 10(7), e0120644. 10.1371/journal.pone.0120644

20. Ólafsdóttir, H. F., Bush, D., & Barry, C. (2018). The Role of Hippocampal Replay in Memory and Planning. Current Biology, 28(1), R37–R50. 10.1016/j.cub.2017.10.073

21. Oostenveld, R., Fries, P., Maris, E., & Schoffelen, J.-M. (2011). FieldTrip: Open Source Software for Advanced Analysis of MEG, EEG, and Invasive Electrophysiological Data. Computational Intelligence and Neuroscience, 2011(1), 156869. 10.1155/2011/156869

22. Pelli, D. G. (1997). The VideoToolbox software for visual psychophysics: Transforming numbers into movies. Spatial Vision, 10(4), 437–442. 10.1163/156856897X00366

23. Petzka, M., Charest, I., Balanos, G. M., & Staresina, B. P. (2021). Does sleep-dependent consolidation favour weak memories? Cortex; a Journal Devoted to the Study of the Nervous System and Behavior, 134, 65–75. 10.1016/j.cortex.2020.10.005

24. Plog, E., Antov, M. I., Bierwirth, P., Keil, A., & Stockhorst, U. (2022). Phase-Synchronized Stimulus Presentation Augments Contingency Knowledge and Affective Evaluation in a Fear-Conditioning Task. *eNeuro*, 9(1). 10.1523/ENEURO.0538-20.2021

25. Plog, E., Antov, M. I., Bierwirth, P., & Stockhorst, U. (2023). Effects of phase synchronization and frequency specificity in the encoding of conditioned fear–a web-based fear conditioning study. PLOS ONE, 18(3), e0281644. 10.1371/journal.pone.0281644

26. Rasch, B., & Born, J. (2013). About sleep’s role in memory. *Physiological Reviews*, 93(2), 681– 766. 10.1152/physrev.00032.2012

27. Scoville, W. B., & Milner, B. (2000). Loss of recent memory after bilateral hippocampal lesions. 1957. The Journal of Neuropsychiatry and Clinical Neurosciences, 12(1), 103–113. 10.1176/jnp.12.1.103

28. Serin, F., Wang, D., Davis, M. H., & Henson, R. (2024). Does theta synchronicity of sensory information enhance associative memory? Replicating the theta-induced memory effect. Brain and Neuroscience Advances, 8, 23982128241255798. 10.1177/23982128241255798

29. Singmann, H., Bolker, B., Westfall, J., Aust, F., & Ben-Shachar, M. S. (2025). afex: Analysis of Factorial Experiments. 10.32614/CRAN.package.afex

30. Stickgold, R., & Walker, M. P. (2013). Sleep-dependent memory triage: Evolving generalization through selective processing. Nature Neuroscience, *16*(2), 139–145. 10.1038/nn.3303

31. Tadel, F., Baillet, S., Mosher, J. C., Pantazis, D., & Leahy, R. M. (2011). Brainstorm: A User-Friendly Application for MEG/EEG Analysis. Computational Intelligence and Neuroscience, 2011(1), 879716. 10.1155/2011/879716

32. Tirole, M., Duvelle, É., & Bendor, D. (2026). Time, but not reward, shapes replay-based episodic prioritization (p. 2026.06.10.731494). bioRxiv. 10.64898/2026.06.10.731494

33. Tononi, G., & Cirelli, C. (2006). Sleep function and synaptic homeostasis. Sleep Medicine Reviews, 10(1), 49–62. 10.1016/j.smrv.2005.05.002

34. Tononi, G., & Cirelli, C. (2014). Sleep and the price of plasticity: From synaptic and cellular homeostasis to memory consolidation and integration. Neuron, 81(1), 12–34. 10.1016/j.neuron.2013.12.025

35. van der Plas, M., Wang, D., Brittain, J.-S., & Hanslmayr, S. (2020). Investigating the role of phase-synchrony during encoding of episodic memories using electrical stimulation. Cortex; a Journal Devoted to the Study of the Nervous System and Behavior, 133, 37–47. 10.1016/j.cortex.2020.09.006

36. Van Veen, B. D., Van Drongelen, W., Yuchtman, M., & Suzuki, A. (1997). Localization of brain electrical activity via linearly constrained minimum variance spatial filtering. IEEE Transactions on Biomedical Engineering, 44(9), 867–880. IEEE Transactions on Biomedical Engineering. 10.1109/10.623056

37. Wang, D., Clouter, A., Chen, Q., Shapiro, K. L., & Hanslmayr, S. (2018). Single-trial phase entrainment of theta oscillations in sensory regions predicts human associative memory performance. Journal of Neuroscience, 38(28), 6299–6309. 10.1523/JNEUROSCI.0349-18.2018

38. Wang, D., Marcantoni, E., Shapiro, K. L., & Hanslmayr, S. (2026). Pre-stimulus alpha power modulates trial-by-trial variability in theta rhythmic multisensory entrainment strength and theta-induced memory effect. Communications Psychology, 4(1), 40. 10.1038/s44271-026-00406-x

